# Modes of migration and multilevel selection in evolutionary multiplayer games

**DOI:** 10.1101/029470

**Authors:** Yuriy Pichugin, Chaitanya S. Gokhale, Julián Garcia, Arne Traulsen, Paul B. Rainey

**Author notes:** Corresponding author: *Email address:* (Yuriy Pichugin).

## Abstract

The evolution of cooperation in group-structured populations has received much attention, but little is known about the effects of different modes of migration of individuals between groups. Here, we have incorporated four different modes of migration that differ in the degree of coordination among the individuals. For each mode of migration, we identify the set of multiplayer games in which the cooperative strategy has higher fixation probability than defection. The comparison shows that the set of games under which cooperation may evolve generally expands depending upon the degree of coordination among the migrating individuals. Weak altruism can evolve under all modes of individual migration, provided that the benefit to cost ratio is high enough. Strong altruism, however, evolves only if the mode of migration involves coordination of individual actions. Depending upon the migration frequency and degree of coordination among individuals, conditions that allow selection to work at the level of groups can be established.

## 1. Introduction

Cooperation can be defined as “a joint action for mutual benefit” (16; 43; 10; 62). Participation in a cooperative act is generally costly to cooperators (27; 6; 10). Therefore, cooperators have lower fitness than non-cooperators (defectors) and, thus, should be eliminated by natural selection. Nevertheless, cooperation is widespread in nature (11; 53; 74). How cooperation evolves and is maintained in the face of selfishness has been the subject of intensive investigation (27; 71; 6; 47; 67).

In a group-structured population, members of cooperative groups have a selective advantage over the members of non-cooperative groups. This advantage can make the evolution of cooperation possible (28; 71; 65; 47). The essential idea is that population structure channels cooperation preferentially to other cooperators (20; 19). Wilson and Wilson (73) formulated this as: “Selfishness beats altruism within groups. Altruistic groups beat selfish groups. Everything else is commentary.” However, the interplay between these effects is important because it determines whether cooperation will evolve.

Group structure by itself does not provide an advantage to cooperation (23) - indeed within groups, selfish types have an advantage over cooperating types (71). For cooperating types to be maintained, groups must participate in some kind of birth and death process. Individuals arising within one group must have an opportunity to become a member of another group. There are many ways by which this may occur. For instance, in standard trait group models (71; 5; 22), individuals within groups are released into a global pool and then randomly form new groups. Alternatively groups may fragment (65). A further possibility is that individuals from one group may migrate to another (9; 37; 31; 32). Via the process of migration, groups themselves do not reproduce in a conventional sense, but the effects are parallel.

In this study we consider models where an individual may become a member of another group by migration between groups. Individuals migrating from one group to another may fixate in the new group, or be eradicated as a consequence of individual-level selection. A defecting individual has a higher probability of fixation in a group of cooperators than does a cooperating individual in a group of defectors, thus individual-level selection favours defectors. However, individuals in groups of cooperators are more productive than in groups of defectors, and therefore groups of cooperators release more migrants than do groups of defectors. Thus, while previous studies have shown that migration makes cooperation more difficult to evolve (because it brings about the mixing of groups (65)), recent work shows that rare migration can favor cooperation (32). Here, we consider a range of modes by which migration might occur and describe ensuing effects on the evolution of cooperation.

Migration can be implemented in multiple ways: individuals may migrate individually, or in clumps; subsequent migrations may or may not be influenced by previous ones; migration may be triggered by signals perceived by individuals, or may be influenced by the group. In this study we compare different modes of migration. For each mode, we identify the games in which cooperation is evolutionarily successful, i.e., where selection at the group level is strong enough to overcome selection at the individual level. The comparison between modes of migration shows that the set of games in which cooperation evolves generally expands with increasing degrees of coordination surrounding the migration process.

## 2. Evolutionary dynamics within a single group

We make the assumption that individuals live in a population with a fixed number of groups. The interactions between all individuals within a group are determined by a multiplayer game. The payoff of each individual depends on its strategy and the composition of the group. Each individual can be either a cooperator (*C*) or a defector (*D*). The size of the game is equal to group size. Thus, all players sharing the same strategy within a group have the same payoff. More specifically, the payoff of a cooperator in a group with *i* cooperators and *n – i* defectors is *a_i_,* and the payoff of a defector in a group with *i* cooperators and *n* – *i* defectors is *b_i_*. Thus, a game is completely determined by two sequences, *a*_1_, &, *a_n_,* and *b*_0_, &, *b_n_*_-1_ (38; 25).

We use an exponential function to map payoff to fitness. The fitnesses of cooperators and defectors in a group with *i* cooperators are therefore 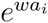, and 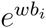, respectively (66). Here, *w* measures the intensity of selection. For *w* = 0, selection is neutral. For *w* ≪ 1, the fitness is approximately linear in payoffs. For large *w*, small differences in payoffs lead to large fitness differences.

The evolutionary dynamics are governed by a Moran process. At each time step a single individual in the population is chosen for reproduction with probability proportional to fitness (45; 48). This chosen individual produces identical offspring, replacing a randomly chosen individual. Thus, population size is kept constant. For such a process, the probability for a single cooperator to take over the whole population, *ϕ_C_*, can be calculated exactly, as well as the probability of a single defector taking over the whole population, *ϕ_D_* (24; 64). These fixation probabilities form the basis of our measure of success for each strategy.

In order to compare the evolutionary success of the two strategies *C* and *D*, we examine whether *ϕ_C_* > *ϕ_D_*. Thus, the value of *ϕ_C_*/*ϕ_D_* determines which strategy is more common. For a ratio greater than 1, cooperation is favoured over defection. If the ratio is less than 1, defection is favoured. The fixation probabilities of cooperators and defectors in the Moran process with exponential mapping are (36; 48; 66)

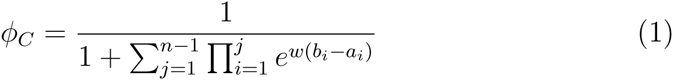

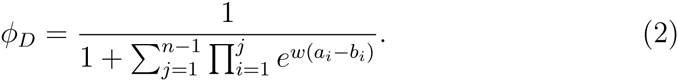

The ratio of the fixation probabilities is given by (46)

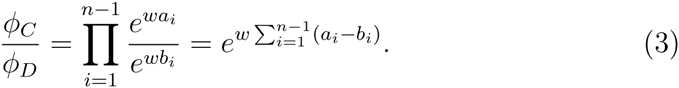

Whether this ratio is greater than 1 (i.e. cooperators are favoured) depends solely on the sign of

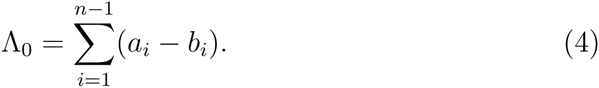

This is a generalization of the classic result of risk dominance to multiplayer games (35; 48; 21; 3; 40; 26). For a positive Λ_0_, cooperation is favoured in terms of the fixation probability, while a negative Λ_0_ means that defectors are selected. We will use such Λ values for comparing the different migration modes.

## 3. Migration modes

We now extend this analysis to multiple groups, and include migration between groups (see Fig. 1). Consider *m* different groups, each with a fixed group size of *n*. We discuss several different modes of migration that individuals can use to move between groups.

**Figure 1:**
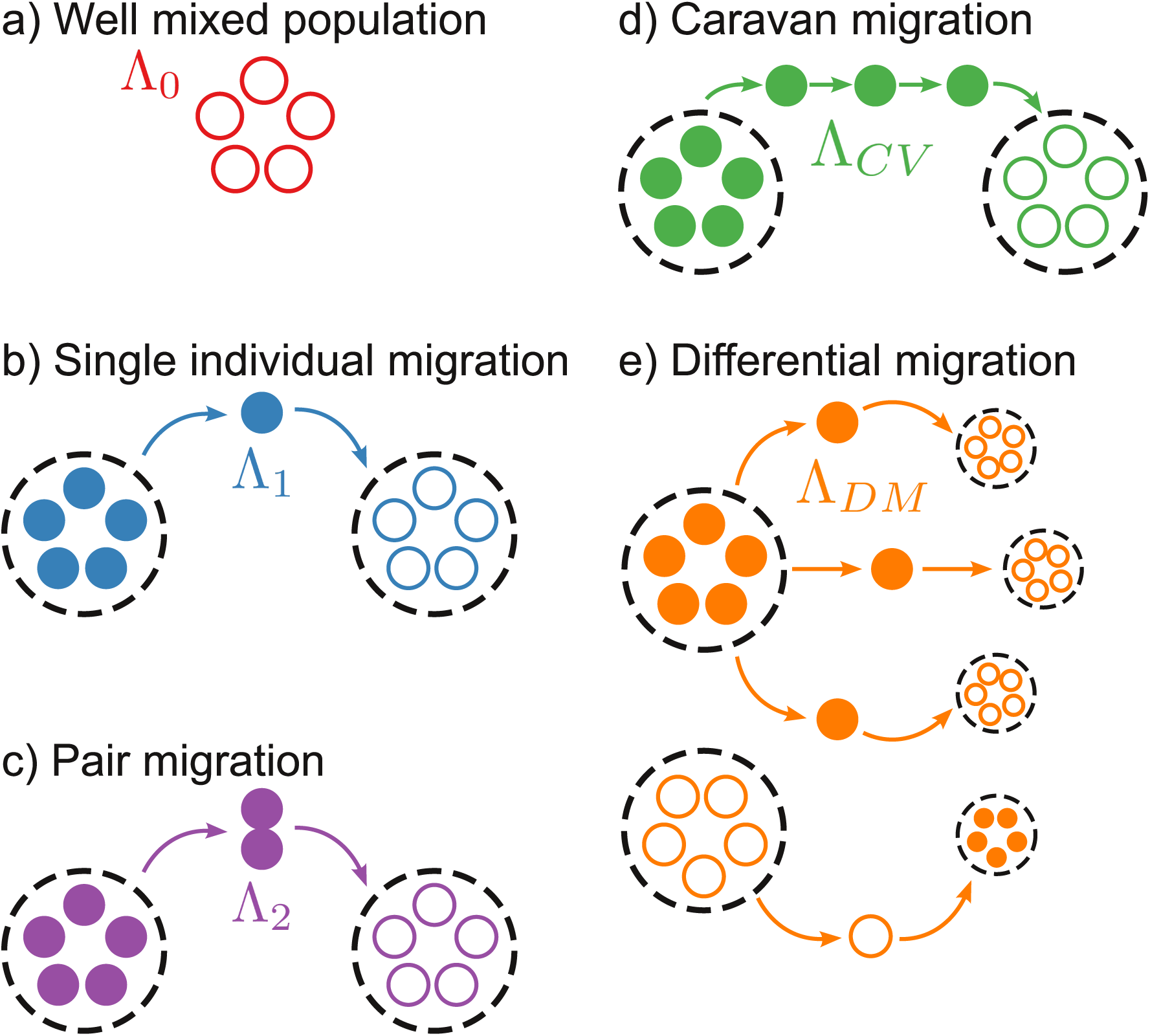
Different modes of migration. Closed circles represent cooperators, open circles represent defectors, dotted line circles represent groups. (a) *Well mixed population,* where no migration is possible. (b) *Single individual migration mode,* where each individual migrates independently. (c) *Pair migration mode,* where individuals migrate in pairs. (d) *Caravan migration mode,* where multiple migrants go to the same group. (e) *Differential migration mode*, where cooperators have higher chances to migrate than defectors. In each case, the quantity Λ determines whether cooperation evolves or not, cf. Fig. 2.

The rate of migration between groups is assumed to be very small compared to the rate of fixation of a strategy within a group. This implies that migration events typically occur only when groups are homogeneous (65; 66). Under this time-scale separation, fixation events in the whole population occur in two stages: first a strategy fixes inside a group – with probability *ϕ_C_* (*ϕ_D_*) for cooperators (defectors) – and then in the whole population – with probability Φ*_C_* (Φ*_D_*) for groups of cooperators (defectors).

We use Eqs. (1) and (2) to compute *ϕ_C_* and *ϕ_D_* at the individual level. At the group level, the fixation probabilities Φ*_C_* and Φ*_D_* depend on the mode of migration. Expressions for these probabilities are generally simpler than for the probabilities at the individual level due to the fact that all individuals within a group have the same fitness when migration occurs (see Appendix A.2-Appendix A.5 for details).

The ratio of fixation probabilities in the structured population (analogous to Eq. (3)) is then given by 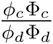 (65).

Here we present a brief derivation of fixation probabilities and corresponding “sign sums” Λ. A different Λ will be calculated for each migration mode (See Appendix A.1 – Appendix A.5 for details).

### 3.1. Single individual migration

As in Traulsen and Nowak (65), we assume that offspring are added to the parent group with probability 1 − *λ*, or to a randomly chosen group with probability *λ*. This is the simplest migration process, with *λ* being the migration probability. Due to *λ* ≪ 1, we consider the probability that a group where the mutant has fixed will send out a migrant that will become a member another group. This probability is equal to 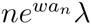 for groups of cooperators, and 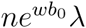 for groups of defectors. For the fixation probabilities at the group level, we obtain the ratio

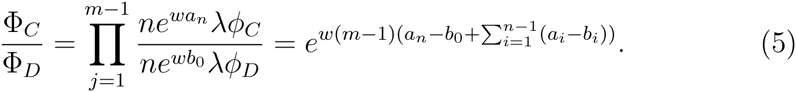

Combining Eqs. (3) and (5) we obtain

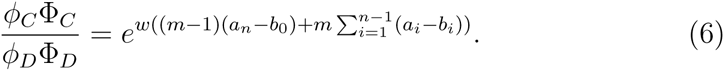

Here, the outcome of evolution is determined by the sign of Λ_1_, given by

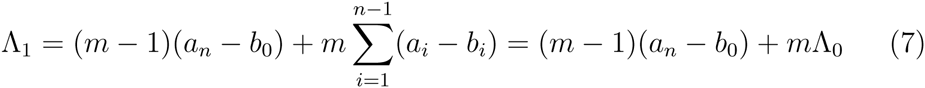

The equation for the sign sum Λ_1_ contains the sign sum of the single group mode, Λ_0_, as the second term. The first term (*m* − 1)(*a_n_* − *b*_0_) is proportional to the fitness difference of the purely cooperative group and the purely defecting group, and describes the effect of group migration. Eq. 7 explicitly expresses conditions for selection to favor cooperation in the single individual migration mode (31) through payoffs from a multiplayer game that is played within groups.

Groups of cooperators send out more migrants than groups containing high frequencies of defecting types, which means that cooperative strategies gain an advantage in the face of migration. The effect of migration depends on the number of groups *m* in a population. The relative weight of the new term in comparison with the lower-level selection term 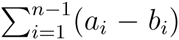 depends only weakly on the number of groups *m*. With decreasing number of the groups, Λ_1_ approaches Λ_0_, and for *m* = 1 both are identical.

### 3.2. Pair migration

Another mode of migration is one where migrants leave simultaneously. For this mode we assume that every migration event carries propagules of a finite number. For illustrative purposes, we discuss propagules of size 2 or ‘pair migration’. In this case, we consider the probability that two deviating individuals take over the population. The sign sum is

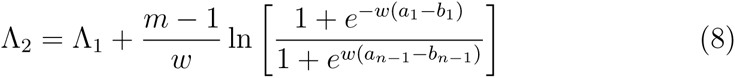

The additional term, now including the selection coefficient, may be positive or negative, depending on the payoff comparison in groups with 1 and *n* − 1 individuals of each type. For a game where cooperators receive a lower payoff than defectors in the same group, this additional term is always positive. Therefore the increase of invading propagule size from 1 to 2 benefits cooperators. The sign sums can be calculated for propagules of arbitrary size in a similar fashion.

### 3.3. Caravan migration

Next, we assume that a new migrant might follow a previous migrant with a probability *p*. This causes a caravan effect, whereby migrants invade the same group with a probability greater than random. Due to the time scale separation assumption, a migrant is fixed or eliminated from the group before the next migrant arrives. Therefore, the caravan migration mode considers multiple migrations of single individuals, whereas the propagule mode of migration considers simultaneous migration of multiple individuals. For simplicity we introduce an additional time scale separation to the caravan migration model: all follow-up migrants arrive at the recipient group earlier than migrants from any other group. The caravan migration mode represents biological systems in which migrants may leave some record of their migration that stimulates the production of further individuals within the group to follow the first departed migrant. This approximates a situation where, for example, an ant leaves a chemical trail (33). This is the simplest example of the model in which all players in the group coordinate their actions.

The probability that the number of migrants entering the same group is equal to *k* is given by

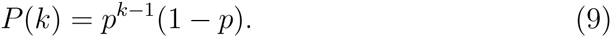

The probability that at least one migrant is successful is equal to

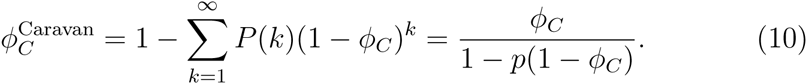

Here 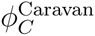 is the probability of a successful invasion of a group of defectors by a cooperative group.

Similarly, the expected probability of the opposite event is 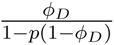. Thus, the ratio of fixation probabilities at the group level is

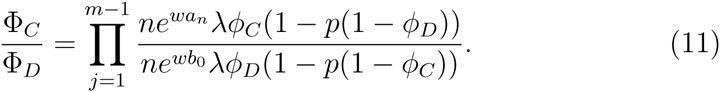

If *p* ≫ 1 − *ϕ*, the probability that the group invaded by the first migrant is eventually taken over approaches 1, such that the result becomes independent of *ϕ_C_* and *ϕ_D_*. The group that receives the first migrant will be invaded with a probability equal to 1. The flow of migrants from one group to another means that the invaded group will be converted with a probability equal to 1. The ratio of fixation probabilities at the group level in this limit is

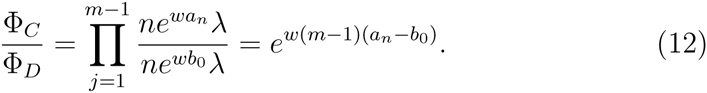

The sign sum (see Eq. (7)) for this mode is then

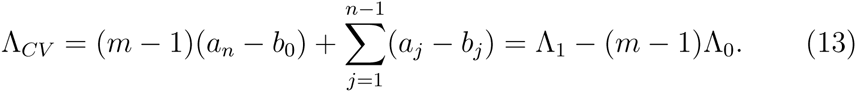

This is larger than in the migration mode for a single individual Λ_1_, as Λ_0_ < 0 for traits that are disadvantageous at the individual level (*a_i_* < *b_i_*). An increase in the number of groups in a population significantly increases the advantage to cooperators caused by this migration process. Since the caravan mode effectively displaces the accepting group with a copy of the donor group, the result obtained here is mathematically equivalent to those of Traulsen and Nowak (65), where it was assumed that a group splits and displaces a randomly selected group.

### 3.4. Differential migration

In the earlier migration modes we have assumed that the migration rate is independent of the type of emigrant. Here we relax this assumption. For example, a group of cooperators may increase the migration rate of its members, therefore increasing the fitness of the group as a whole. Biologically, this could be envisioned to occur via secretion of a chemical signal promoting newly emerged individuals to leave the parent group.

In this mode, let *λ_C_* be the migration rate of *C* types, and *λ_D_* be the migration rate of *D* types. Assuming that the time scale separation is not violated by increased migration rates, we calculate the ratio of fixation probabilities on the group level as

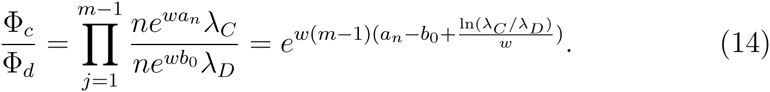

Therefore, the sign sum in this mode is

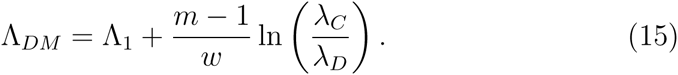

The difference in migration rates (*λ_C_* > *λ_D_*) provides an advantage to cooperating groups, which emits proportionally more migrants in this mode. This is reflected in an additional term 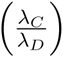, which can shift the balance of selection in favour of cooperators. Interestingly, the overall sign sum Λ*_DM_* may still be negative, despite the fact that groups of cooperators produce more migrant offspring than groups of defectors. This can be explained by the fact that the raw number of migrant offspring is not a determinant of evolutionary success, instead the number of successfully invaded migrants is a defining characteristic of evolution in our model. As such, even if the number of migrants emitted by the cooperating group might be high, the fixation process occurring by means of selection at the individual level favours defectors. The interplay of these two factors does not necessarily promote cooperation even in the differential migration mode, where cooperators are considered to have an advantage.

## 4. Social dilemmas

To be more concrete, we now apply the results of the previous sections to different social dilemma games (12; 6; 38; 47). In social dilemma games, the average payoff to players increases with the number of cooperators, but defectors gain higher payoff than cooperators. An example of a pairwise social dilemma is the prisoner’s dilemma, which is extensively used for the study of the evolution of cooperation (6; 44; 15). For our purposes, it is useful to differentiate between weak and strong altruism.

### 4.1. Weak and strong altruism

Weak altruism is a situation where cooperators provide an advantage to the group, but regardless of the group composition, cooperators have lower payoff than defectors (72; 38). Therefore, the payoffs under weakly altruistic interactions have two properties,

1. If the number of cooperative players increases, the payoffs of all players increase. That is *a_i_* < *a_i_*_+1_ and *b_i_* < *b_i_*_+1_.
2. Cooperators have lower payoff than defectors. That is *a_i_* < *b_i_*.

Since *a_i_* < *b_i_*, then, consequently, Λ_0_ < 0. Unsurprisingly, weak altruism does not arise in the absence of selection at the group level.

In the case of single migrants, the migration-related term in the sign sum Λ_1_ (Eq. (7)) can balance, and even overcome, the term that represents lower level selection. Thus, weak altruism can be favoured in simple migration settings. Similar arguments hold for the pair migration, caravan migration and differential migration modes.

Strong altruism (72), also termed as focal complement altruism (38), are interactions where switching to cooperation always entails a loss of reproductive success. A well-known example of strong altruism is the prisoner’s dilemma where strongly altruistic interactions are characterized by two properties:

1. If the number of cooperative players increases, the payoffs of all players increase. That is *a_i_* < *a_i_*_+1_ and *b_i_* < *b_i_*_+1_.
2. If a player switches from defection to cooperation, their payoff decreases. That is *a_i_* < *b_i_*_—1_.

Strong altruism is always disadvantageous in populations without structure, i.e. Λ_0_ < 0. In addition, we find that

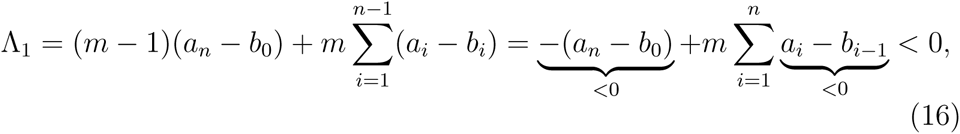
which means that strong altruism is also disfavoured with simple individual-based migration. This result generalizes previous findings that cooperation in the Prisoner’s dilemma game cannot evolve when migration involves just a single individual (31).

For pair migration, Λ_2_ can become positive due to the additional term that is present in Λ_2_ (see Eq. 8). Also for caravan migration, cooperation can be favored due to the additional positive term – (*m* – 1)Λ_0_.

Next, we discuss more specific examples of social dilemmas.

### 4.2. Public goods games

Pairwise games, such as the prisoner’s dilemma, where only two players participate in each game round, cannot represent cooperation with synergistic interactions. With synergistic interactions, multiple cooperators amplify each other’s contributions, thus providing higher benefit than they would produce independently. To encompass these kinds of interactions, we utilize *multiplayer games,* where multiple players are taken into account in the payoff calculation (30; 40; 26).

Public goods games are a type of multiplayer game where each player can make a donation to a public pool. The collected amount is then multiplied, and evenly shared amongst all players, including those that decided not to make a donation. Weak and strong altruism can be naturally represented by self-returning benefit and self-excluding benefit games, respectively (59; 14). In self-returning benefit games, the public goods are shared among all participants; therefore, a proportional part of a donation returns to contributors as a part of their payoff. In this case, all players receive the same share of a public good, but defectors save the cost of donation. Therefore self-returning benefit games represent weak altruism. In self-excluding benefit games, a donation by a focal individual is only shared among other participants; therefore, the payoff of this focal player depends only on the donation of others. In self-excluding benefit games, switching from cooperation to defection does not change the received amount of the public goods, but saves the cost of cooperation. This makes cooperation in self-excluding benefit games strongly altruistic.

We start with the simplest public goods game. Here, the reward to cooperators increases linearly with the number of cooperators. Cooperative individuals pay a cost *γ*, in order to provide a benefit *β*. This benefit is either split amongst the rest of the group, in the linear self-excluding game (LSE game); or split among the whole group, in the linear self-returning game (LSR game). A defecting individual does not pay the cost, but reaps the benefits from other cooperators. The LSR game is weakly altruistic, see Table 1. The LSE game is strongly altruistic, and can be viewed as a multiplayer generalization of the standard prisoner’s dilemma.

**Table 1:** Payoffs and their differences for the linear self-returning (LSR), the linear self-excluding (LSE), the synergy/discounting self-returning (SDSR), and the synergy/discounting self-excluding (SDSE) public goods games. *a_i_* is the payoff to a cooperator in a group of size *n* with *i* cooperators, and *b_i_* is the payoff for a defector. The sum of the payoff difference *a_i_* – *b_i_* determines the value of Λ_0_ (see Eq. 4). Switching from defection to cooperation leads to a payoff difference *a_i_* – *b_i_*_-1_. Switching always decreases payoffs for the self-excluding benefit games (LSE and SDSE); however, the change in payoff for the self-returning benefit games (LSR and SDSR) can be positive and therefore cooperators could have a higher fixation rate than defectors in these games.

|  | LSR | LSE | SDSR | SDSE |
| --- | --- | --- | --- | --- |
| $a_i$ | $\frac{i}{n}\beta - \gamma$ | $\frac{i-1}{n-1}\beta - \gamma$ | $\frac{\beta}{n} \frac{1-\zeta^i}{1-\zeta} - \gamma$ | $\frac{\beta}{n-1} \frac{1-\zeta^{i-1}}{1-\zeta} - \gamma$ |
| $b_i$ | $\frac{i}{n}\beta$ | $\frac{i}{n-1}\beta$ | $\frac{\beta}{n} \frac{1-\zeta^i}{1-\zeta}$ | $\frac{\beta}{n-1} \frac{1-\zeta^i}{1-\zeta}$ |
| $a_i - b_i$ | $-\gamma < 0$ | $-\frac{\beta}{n-1} - \gamma < 0$ | $-\gamma < 0$ | $-\frac{\beta}{n-1}\zeta^{i-1} - \gamma < 0$ |
| $a_i - b_{i-1}$ | $\frac{\beta}{n} - \gamma$ | $-\gamma < 0$ | $\frac{\beta}{n}\zeta^{i-1} - \gamma$ | $-\gamma < 0$ |
| Altruism | Weak | Strong | Weak | Strong |

In addition, we consider non-linear public goods games. If there are synergies in the production of the public goods, each additional donation can provide more benefits than the previous one. Likewise, if the marginal benefit decreases with the number of donations, the benefits are discounted and become saturated as the number of cooperators increase. These so-called non-linear public goods games have been extensively analyzed (18; 7; 30; 69; 50; 25; 70; 4; 51; 55; 1).

In the simplest version of the game incorporating synergy and discounting, the first cooperator in the group pays a cost *γ* to generate *β* units of a public good. Each additional cooperator present in the group provides *ζ* times the public good than the previous one. If *ζ* > 1, then cooperators act synergistically. If *ζ* < 1, benefits are discounted. Again, donations can be either shared among all players (synergy/discounting game with self-returning benefit, or SDSR game), or only among other players and excluding the donor (synergy/discounting game with self-excluding benefit, or SDSE game).

The payoffs *a_i_* and *b_i_* to players in these games (LSR, LSE, SDSR and SDSE) and their differences are presented in Table 1. For each of these games we derive the conditions for the evolution of cooperation under different migration schemes (see Section 3). The sign sums for each combination of game and migration mode are presented in Tables 2 (for self-returning games) and 3 (for self-excluding games).

**Table 2:** Sign sums for self-returning games (weak altruism) in a well mixed population and under different modes of migration.

|  | LSR | SDSR |
| --- | --- | --- |
| $\Lambda_0$ | $-(n-1)\gamma$ | $-(n-1)\gamma$ |
| $\Lambda_1$ | $(m-1)\beta - (mn-1)\gamma$ | $(m-1)\frac{\beta}{n}\frac{1-\zeta^n}{1-\zeta} - (mn-1)\gamma$ |
| $\Lambda_2$ | $(m-1)\beta - m(n-1)\gamma$ | $(m-1)\frac{\beta}{n}\frac{1-\zeta^n}{1-\zeta} - m(n-1)\gamma$ |
| $\Lambda_{CV}$ | $(m-1)\beta - (m+n-2)\gamma$ | $(m-1)\frac{\beta}{n}\frac{1-\zeta^n}{1-\zeta} - (m+n-2)\gamma$ |
| $\Lambda_{DM}$ | $(m-1)\left(\beta + \frac{\ln(\lambda_C/\lambda_D)}{w}\right) - (mn-1)\gamma$ | $(m-1)\left(\frac{\beta}{n}\frac{1-\zeta^n}{1-\zeta} + \frac{\ln(\lambda_C/\lambda_D)}{w}\right) - (mn-1)\gamma$ |

**Table 3:** Sign sums for self-excluding public good games (strong altruism) in a well mixed population and under different modes of migration.

|  | LSE | SDSE |
| --- | --- | --- |
| $\Lambda_0$ | $-\beta - (n-1)\gamma$ | $-\frac{\beta}{n-1} \frac{1-\zeta^{n-1}}{1-\zeta} - (n-1)\gamma$ |
| $\Lambda_1$ | $-\beta - (mn-1)\gamma$ | $-\frac{\beta}{n-1} \frac{1-\zeta^{n-1}}{1-\zeta} - (mn-1)\gamma$ |
| $\Lambda_2$ | $\beta \frac{m-n}{n-1} - m(n-1)\gamma$ | $-\frac{\beta}{n-1} \frac{1-\zeta^{n-1}}{1-\zeta} - (mn-1)\gamma + \frac{m-1}{w} \ln \left[ \frac{1+e^{w(\frac{\beta}{n-1}+\gamma)}}{1+e^{-w(\frac{\beta}{n-1}\zeta^{n-2}+\gamma)}} \right]$ |
| $\Lambda_{CV}$ | $(m-2)\beta - (m+n-2)\gamma$ | $(m-2) \frac{\beta}{n-1} \frac{1-\zeta^{n-1}}{1-\zeta} - (m+n-2)\gamma$ |
| $\Lambda_{DM}$ | $(m-1) \frac{\ln(\lambda_C/\lambda_D)}{w} - \beta - (mn-1)\gamma$ | $(m-1) \frac{\ln(\lambda_C/\lambda_D)}{w} - \frac{\beta}{n-1} \frac{1-\zeta^{n-1}}{1-\zeta} - (mn-1)\gamma$ |

Sign sums as functions of benefit *β* for different games and modes of migration are presented in Fig. 2. In the well mixed model, cooperation is evolutionary unsuccessful in all games (Λ_0_ < 0).

**Figure 2:**
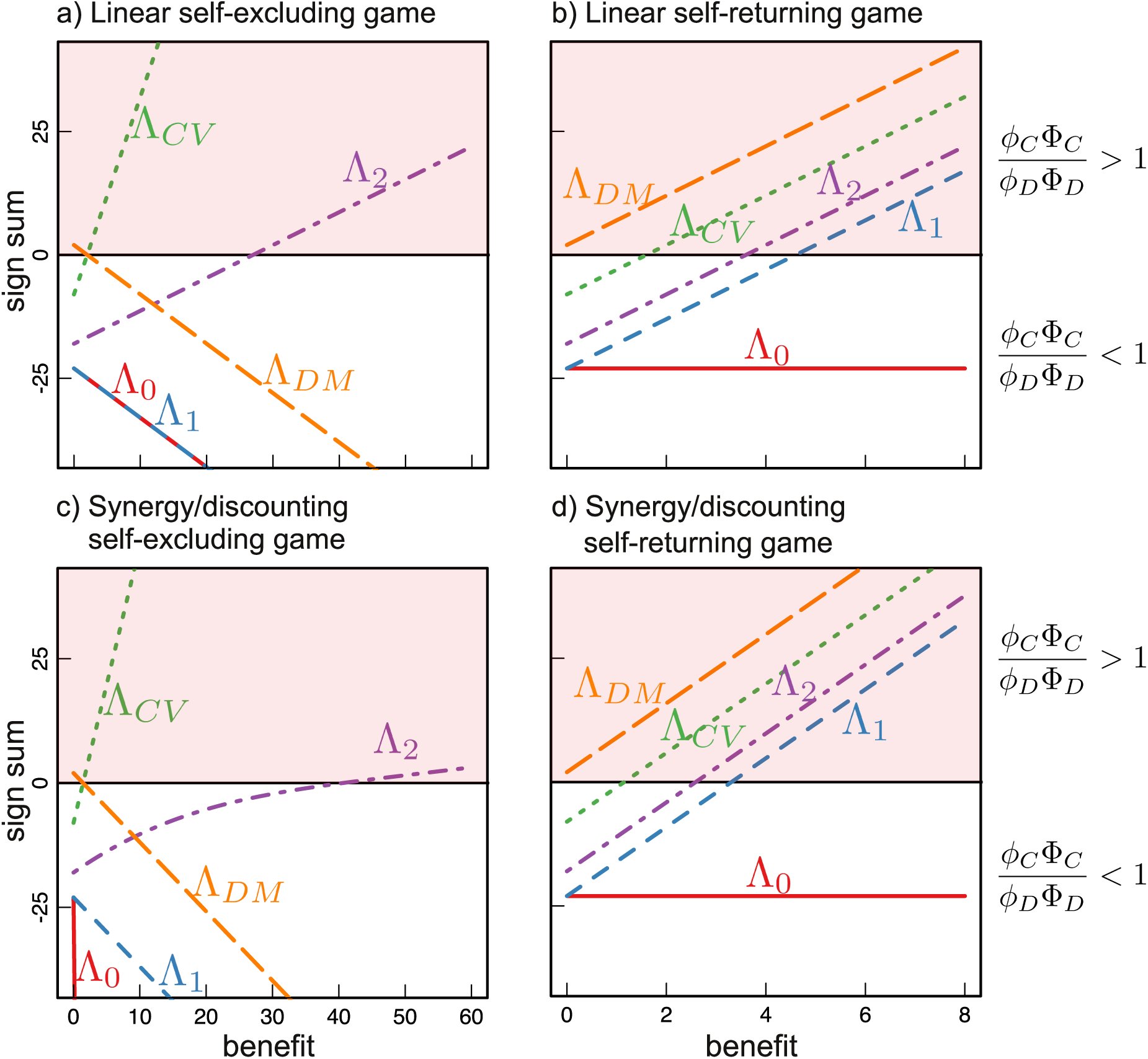
The evolution of cooperation does not always become easier with increasing benefit *β*. Cooperation is advantageous in terms of fixation probabilities if the sign sum Λ calculated for migration modes (lines) is positive (shaded region). In the well mixed case, Λ_0_ decreases in self-excluding games and stays constant for self-returning games. In the single individual migration mode and the differential migration mode, cooperation becomes easier with increasing benefit in the self-returning case, but harder in the self-excluding case. In the pair migration mode and in caravan migration, cooperation becomes easier with increasing benefit for all games with the current parameter set. (*n* = 24 for the well mixed population, *m* 14 = 6 and *n* = 4 in migration models, *ζ* = 1.35, intensity of selection *w* = 0.1, cost of cooperation *γ* = 1, group migration bonus factor in differential migration mode *w*^−1^ ln(*λ_C_/λ_D_*) = 5, colors as in Fig. 1).

For the games representing weak altruism (LSR and SDSR), cooperation may be successful in all migration modes, provided that the benefit to cost ratio is large enough. Clearly, increasing synergy in self-returning games favours cooperation.

For the games representing strong altruism (LSE and SDSE), even for the individual migration mode, cooperators have no selective advantage (Λ_1_ < 0). For the pair migration and caravan migration modes, strong altruism may have a selective advantage over defection and in LSE game this is possible if the benefit to cost ratio is high enough. Under differential migration strong altruism also can have a selective advantage in the LSE game. However, the prerequisites for this are restrictive: the group migration bonus factor *w*^−1^ ln(*λ_C_*/λ*_D_*) must be high enough to ensure a strong implicit advantage to cooperators. Interestingly, an increase in the benefit to cost ratio works against cooperation under this mode of migration.

At the qualitative level, the difference between linear and non-linear games from the same migration scheme are minor, with a few notable exceptions. Strongly altruistic, non-linear SDSE games, can promote cooperation in the pair migration mode at high values of benefit *β* (the sign sum in this case cannot be reduced to benefit to cost ratio) only if the number of groups *m* is high enough (Appendix A.6). The minimal number of groups necessary for the success of cooperation for this game increases with the increase of the synergy. Therefore synergistic interactions work against cooperation success in the pair migration mode.

In the caravan migration mode, the SDSE game, similar to the linear LSE game, promotes cooperation if the benefit to cost ratio (*β*/*γ*) is high enough. However, synergy of cooperators makes cooperation successful at lower values of benefit to cost ratio than in the LSE game. Finally, in the SDSE game with differential migration, as well as in the LSE game, the advantage of cooperation depends on the group migration bonus factor, while both high benefit to cost ratio and synergy work against cooperation.

Synergy always favours cooperation in weakly altruistic self-returning games (LSR and SDSR); however, it may work against cooperation in strongly altruistic LSE and SDSE games under certain modes of migration. Intuitively, cooperation will be enhanced if the benefit provided by a cooperator is large or if there is more synergy between cooperators (larger *β*, larger or increasing *ζ*). Counterintuitively, in self-excluding games this works against cooperation (49). Consider the prisoner’s dilemma game, played by one cooperator and one defector. An increase in the amount of benefit produced by cooperator *β* leads only to an increase in the payoff to the defector; thereby harming cooperation. Furthermore, in a multiplayer game, an increasing *ζ* just provides more benefit for defectors to exploit, as it does not return benefit to the contributor. This shows that cheaper cooperation can benefit defectors.

In all four games, for all values of the benefit to cost ratio, defection is favoured (negative sign sums) in well mixed populations. This illustrates that even weak altruism is less successful than defection in the absence of population structure. The standard migration mode allows LSR and SDSR games to have a positive sign sum if the benefit to cost ratio is large enough. However, cooperators in LSE and SDSE games, being strongly altruistic are always disadvantageous, independent of the benefit to cost ratio.

## 5. Discussion

We have shown that migration, even in the absence of coordination between individuals, promotes the evolution of weakly altruistic cooperation. The single individual migration mode presented here is not based on processes that involve an entire group (65), or specific structure of groups (42). Our results indicate that cooperation may emerge by means of group-level selection even if selection is conducted by the non-coordinated actions of individuals. In other words, selection on the group level can be mediated by population structure alone.

In modes where migration involves the coordinated actions of multiple individuals, cooperation can evolve in a much wider range of games than in the single individual migration mode. In the pair, caravan and differential migration modes, strong altruism can be favored. Also, in weakly altruistic games the range of parameters promoting the evolution of cooperation is extended: the domain of benefit to cost ratio with positive sign sums becomes wider than in the single individual migration mode (see Figure 2 panels b and d). Thus, introduction of coordination between individuals’ actions substantially extends the set of conditions under which cooperation may evolve.

Throughout this manuscript, we have concentrated on the exponential payoff to fitness mapping, which allows a very compact representation of the sign sums. However, many of our results hold for more general payoff to fitness mappings (75; 76). For example, for any mapping in which the number of emitted migrants is proportional to the reproductive output of the players within the group the single individual migration mode can favor weak altruism, but not strong altruism, see Appendix B. This is in contrast to a scenario of a pairwise comparison process (31; 32), where production of migrants moving between groups depends directly on payoffs, but the competition between types within the group depends on differences between individual payoffs.

The evolution of cooperation under limited coordination of individuals’ actions may have particular importance for understanding early stages of the evolution of multicellularity. While details remain unclear, there is general agreement that the earliest stages involved the evolution of simple, undifferentiated groups of cooperating cells (68; 57; 52; 2; 39; 29). In theoretical models of the evolution of cooperation, the mechanistic details surrounding the re-distribution of individuals among groups are often overlooked. Two broad kinds of group formation are generally considered: groups originating from growth of a single individual, referred to as “staying together”, and groups formed by aggregation of individuals, referred to as “coming together” (63). An example of the “staying together” mode is fragmentation (65), as found in the algae *Gonium pectorale* (61). The “coming together” mode is utilized by slime molds (8) and in trait group models (71). Multiple individual modes of group formation can be constructed within these two kinds (71; 5; 52; 65; 54; 22; 41; 60; 63), including those in which ‘staying together’ is combined with migration events that establish new groups (56; 41; 58; 29). From a mechanistic point of view, modes of individual assignment, considered in the previous paragraph, are typically assumed to arise by the coordinated actions of multiple individuals in the group. However, early cellular groups were most likely unable to act as a single coordinated unit, and as such recurrence of these groups was presumably conducted by unregulated actions of individual cells (13).

Based on our results, one can perform a classification of multilevel selection models based on the level of complexity of the interactions between groups. The first class consists of models in which processes between groups are mediated by a single individual, such as the single individual migration mode and metapopulation models (17; 34). As shown here, these kinds of models can promote weak altruism, but not strong altruism. The second class of models are those in which between-group processes involve several individuals, or even whole groups. Examples include the pair, caravan and differential migration modes, and also the splitting of whole groups (65). In the context of the early stages of the evolution of multicellularity, the first class of models likely have particular importance, as the multi-individual actions of the second class generally require coordinated activity, which might not be available for early groups.

## 6. Acknowledgements

The work was supported by the Marsden Fund Council from government funding administered by the Royal Society of New Zealand. P.B.R. holds an International Blaise Pascal Research Chair funded by the French State and the Ile-de-France, managed by the Fondation de l’Ecole Normale Supérieure.

## Appendix A. Derivation of sign sums

### Appendix A.1. *Derivation of* Λ_0_

The fixation probability for one individual of the two strategies in a well mixed population is equal to (46; 64)

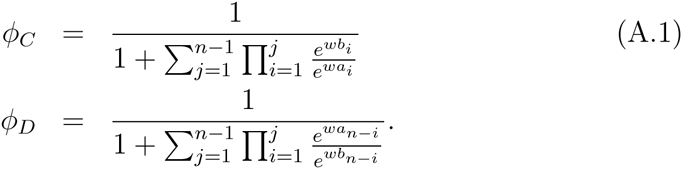

The ratio of these fixation probabilities is

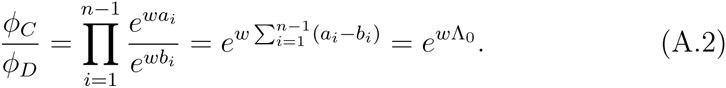

Here, 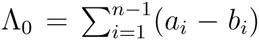 is the sign sum for a well mixed population, as stated previously, for example by (40) and (25).

#### Appendix A.2. *Derivation of* Λ_1_

For group structure and small migration rates, the trait of interest first needs to fix in a group (*ϕ_C_*) and then that group needs to fix in the population (Φ*_C_*). The total fixation probability ratio is thus equal to 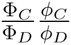 (65). Here 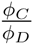 is calculated according to Eq. (A.2). The ratio 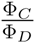 is calculated as

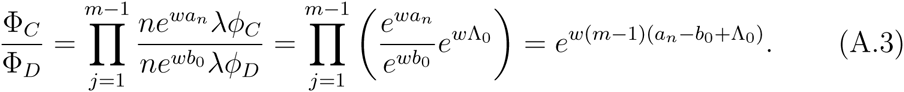

Therefore the total fixation probability ratio is

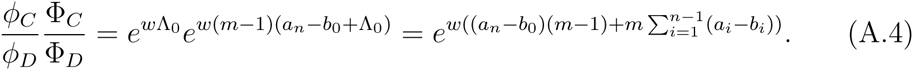

Thus, the sign sum for the single individual migration mode is

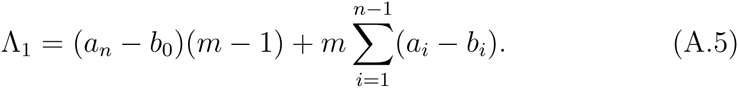

### Appendix A.3. *Derivation of* Λ_2_

In the pair migration mode, the individual-level fixation probabilities are different from the single individual migration mode because the initial state of the group with mixed composition after accepting a migrant is *n*–2 players of the base type and two players of the invading type. Therefore, the fixation probabilities are not equal to the ones presented in Eq. (A.1). According to (46), the fixation probabilities *ϕ^i^* for an initial number of *i* individuals solves the recurrence equation

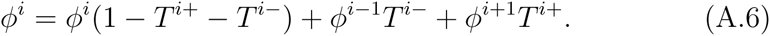

Here *T^i^*^+^ = & and *T^i^*^−^ = & are probabilities to increase or decrease the number of players with a chosen strategy if there are currently i players. Because *ϕ*^0^ = 0, *ϕ*^2^ is

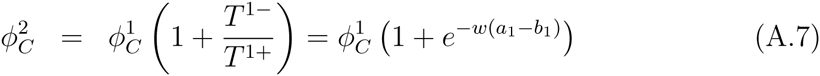

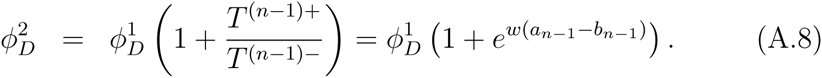

Therefore, the ratio of individual-level fixation probabilities in the pair migration mode is

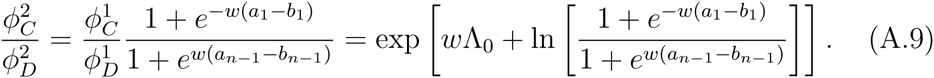

The total ratio of fixation probabilities (taking into account that the invading strategy starts with one player in the first group, and with two players in all following migration invasions) is

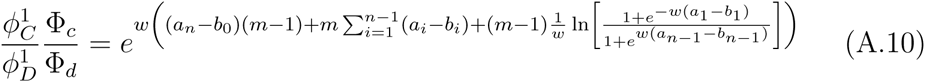

and the sign sum is

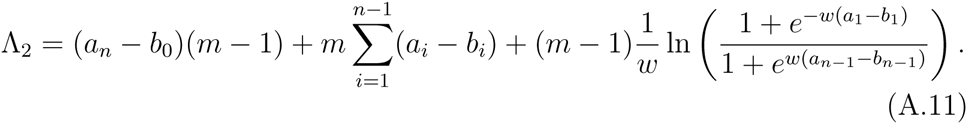

### Appendix A.4. *Derivation of* Λ*_CV_*

In the caravan migration mode with large *p*, the probability of successful invasion of one group into another is equal to 1. Therefore, the ratio of group-level fixation probabilities is

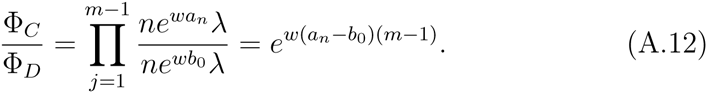

This way the total fixation probabilities ratio is

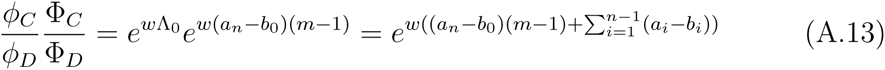

and the sign sum in the caravan migration mode is

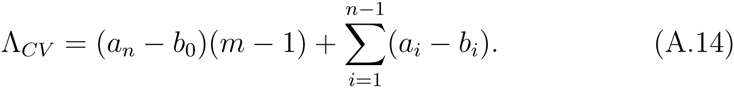

### Appendix A.5. *Derivation of* Λ*_DM_*

In the differential migration mode, the groups have control over the migration probabilities of the players. This affects the fixation probabilities at the group level. The migration probabilities no longer cancel,

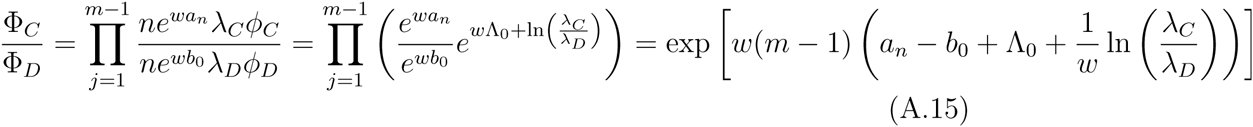

Therefore, the total fixation probability ratio is

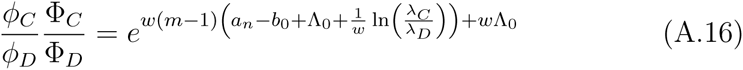

and the sign sum becomes

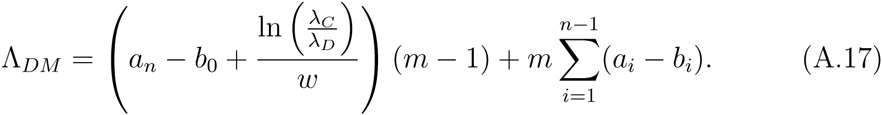

### Appendix A.6. The SDSE game in the pair migration mode

The sign sum for the SDSE game in the pair migration mode is:

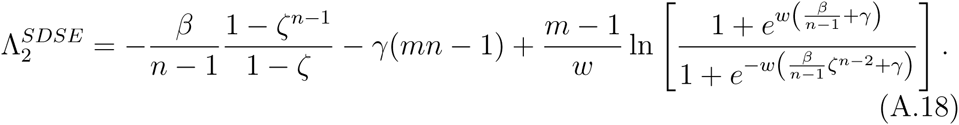

If benefit *β* is high enough 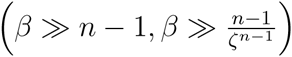, then the sign sum approaches

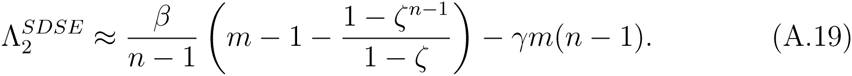

Therefore, the sign sum is positive at high benefit values, if the number of groups *m* is high enough: 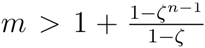. In the case of the discounting game (*ζ* < 1), this condition is more restrictive than *m* ≥ 2, which is always required in multilevel selection models.

## Appendix B. Other payoff to fitness mappings

Strong altruism is at a disadvantage in the single individual migration mode, when we use the exponential payoff to fitness mapping (66). Here we show that this result holds true with any mapping.

In terms of fitness, strong altruism is characterized by two properties:

1. If the number of cooperative players increases, the payoffs of all players increase. That is *f_a_*(*i*) < *f_a_*(*i* + 1) and *f_b_*(*i*) < *f_b_*(*i* + 1), where *f_a_*(*i*) (*f_b_*(*i*)) is the fitness of cooperators (defectors) in a group with *i* cooperators.
2. If a player switches from defection to cooperation, their payoff decreases. That is *f_a_*(*i*) < *f_b_*(*i* – 1).

The ratio of fixation probabilities in the structured population is given by 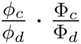 (65). We calculate each term separately.

The ratio of fixation probabilities of a single cell in a group of opposite composition (36; 46) is

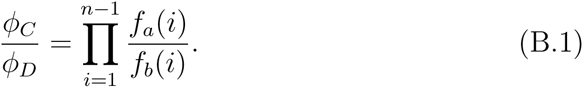

The ratio of fixation probabilities of a single group in a population of opposite composition is

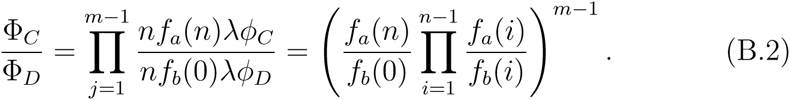

Combining Eqs. (B.1) and (B.2) we get the ratio of fixation probabilities of a single cell in a population of opposite composition:

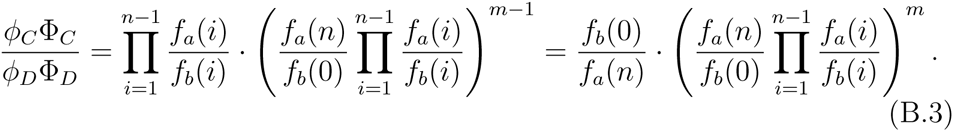

Expression in parenthesis can be rewritten:

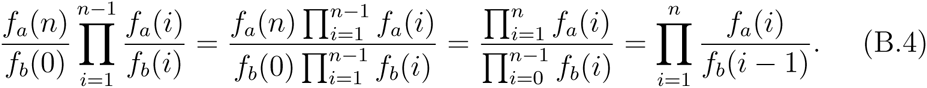

Thus, the fixation probabilities ratio is equal to

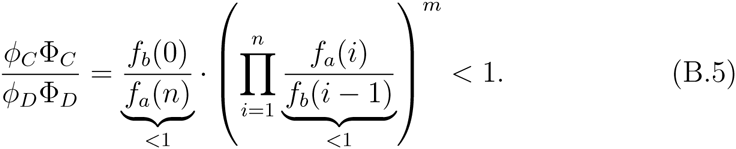

So, the inability of the strong altruism to emerge in a single individual migration mode holds true for all possible payoff to fitness mappings.

